# Rapidly self-sterilizing PPE capable of 99.9% SARS-CoV-2 deactivation in 30 seconds

**DOI:** 10.1101/2020.11.16.384040

**Authors:** Alfred A. Zinn, Mina Izadjoo, Hosan Kim, Kylene Kehn-Hall, Caitlin Woodson, Rachel L. Brody, Robert R. Roth, Agustin Vega, Khanh K. Nguyen, Nhi T. Ngo, Hannah T. Zinn, Lauren Panny, Rafaela Flor, Nicholas Antonopoulos, Randall M. Stoltenberg

**Affiliations:** Kuprion, Inc. San Jose, CA 95134, USA; Integrated Pharma Services, Frederick, MD 21704, USA; Department of Biomedical Sciences and Pathobiology, Virginia-Maryland College of Veterinary Medicine, Virginia Polytechnic Institute and State University, Blacksburg, VA 24061, USA; National Center for Biodefense and Infectious Diseases, School of Systems Biology, George Mason University, Manassas, VA, USA

## Abstract

The coronavirus disease 2019 (COVID-19) has created an acute worldwide demand for sustained broadband pathogen suppression in households, hospitals, and public spaces. The US recently passed a new sad milestone of 500,000 deaths due to COVID-19, the highest rate anywhere in the world. In response, we have created a rapid-acting, self-sterilizing PPE configurations capable of killing SARS-CoV-2 and other microbes in seconds. The highly active material destroys pathogens faster than any conventional copper configuration. The material maintains its antimicrobial efficacy over sustained use and is shelf stable. We have performed rigorous testing in accordance with guidelines from U.S. governing authorities and believe that the material could offer broad spectrum, non-selective defense against most microbes via integration into masks and other protective equipment.

**Summary:** A novel configuration of copper offering continued fast-acting protection against viruses and bacteria, including SARS-CoV-2.

## Introduction

The rush on PPE from the pandemic’s onset illustrated a fatal flaw in our approach to microbe management. The world relies on stockpiles of resources to create a clean field that is sullied immediately upon contact. The sterility provided by widely used disinfectant options is all too fleeting. Demands imposed by COVID-19 turned inefficiencies into breaking points; there is simply not enough sterile equipment in the world to shield ourselves if materials do not offer lasting protection. The masks we are using, while effective, accumulate bacteria over the course of normal wear and are thus limited to a single, short-term use before they must be washed or discarded.^1^ We need a material that keeps hands, surfaces, and tools free of pathogens over long periods of time.

Our engineered copper material (ECM) was first produced as a novel all-copper alternative to tin-based solder. Catalysis taught us that a material’s surface reactivity is contingent on its surface area and energy; the larger the surface area, the more reactive it will be.^2, 3^ ECM owes its activity to a large interconnected meso-structured metal network consisting of a polymeric gel-like copper phase with meso-scale porosity and surface roughness. As COVID-19 overwhelmed the world in early 2020, we hypothesized that this unique structure and its surfactant shell should be highly antimicrobial. The literature is rife with examples of copper as an antimicrobial and the EPA has approved hundreds of copper alloys for their disinfectant properties.^4^–^6^ However, they are slow-acting and require up to 4 hours to fully deactivate microbes. Because conventional copper surfaces disinfect more slowly, they in turn still need to be disinfected regularly. Given the structural differences between ECM and previous copper iterations, it made sense to explore ECM’s anti-COVID-19 activity.

ECM is manufactured using a bottom-up synthesis approach via NaBH_4_ reduction of CuCl_2_ in solution (Fig. 1). The resultant copper material is rendered stable through the formation of a very sticky paste-like metallic gel. It behaves like a typical non-Newtonian “liquid” with high thixotropy (Fig. 5). The specially designed amine surfactant layer exerts strong cohesion via hydrophobic interactions and organic end-chain entanglement, which holds the dense metallic gel together with the aid of the high surface area and energy (Fig. 2) and does not dissolve in water and most organic solvents. Its structure can be described as a dense but flowable metallic gel-like paste with an interconnected meso-scale network, porosity and surface roughness that enables safe handling and processing; it never turns into an airborne powder but rather hardens into a solid copper mass similar to a xerogel, allowing the material to retain the high activity of the unfused ECM.

**Fig 1.**
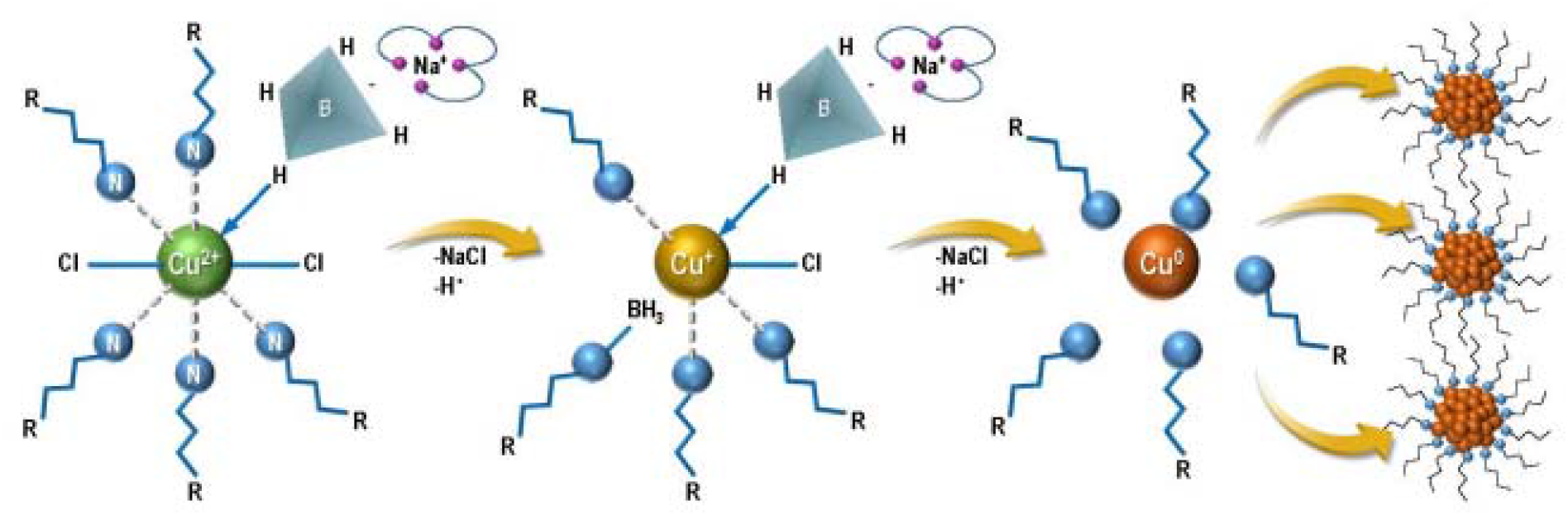
Synthesis of ECM via reduction of CuCl_2_. Anhydrous copper(II) chloride is reduced with sodium borohydride in the presence of amine surfactants to form the raw ECM material. The reaction takes place near room temperature. The borane generated during the reduction forms borane-amine complexes which are later hydrolyzed upon the addition of water to remove NaCl byproducts. The amines assist in dissolving the copper(II) chloride and mediate resultant particle sizes. This prevents the ECM particles from oxidizing and continuing to grow.

**Fig 2:**
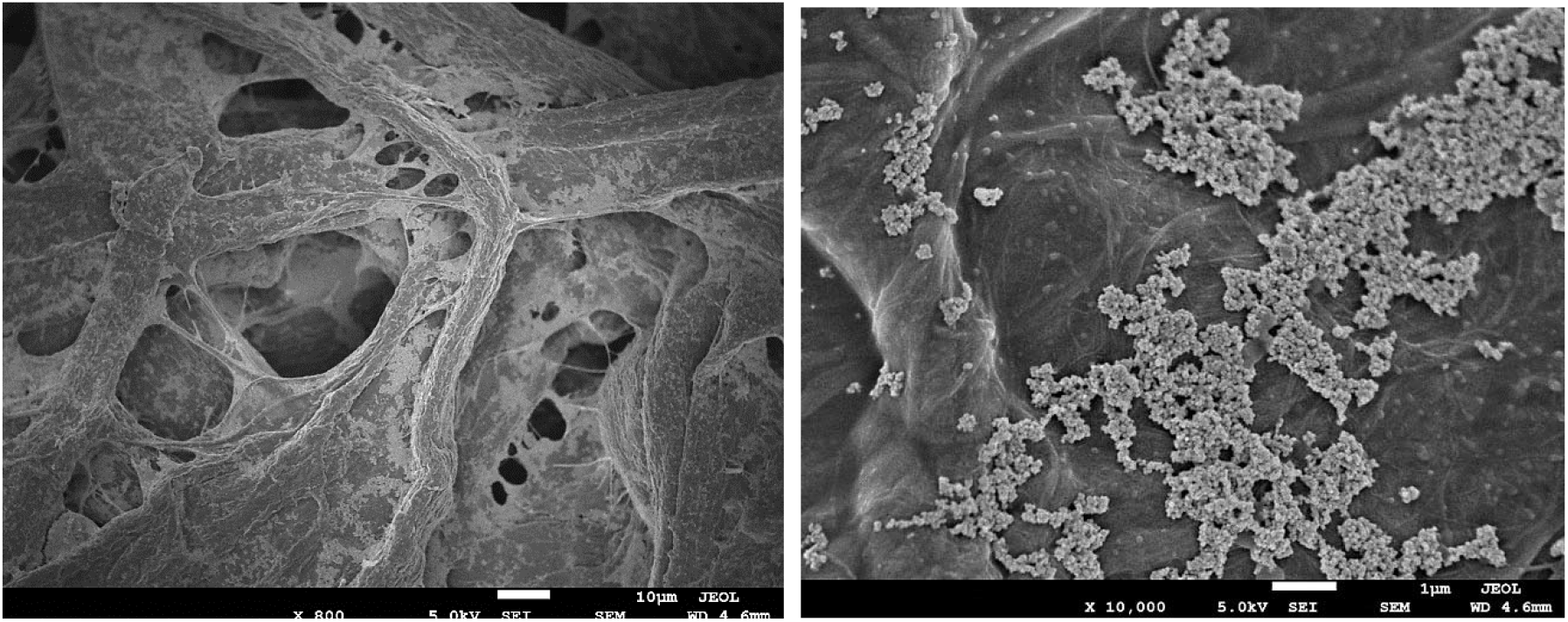
High magnification scanning electron microscopy images of ECM on 45/55 cellulose/polyester fabric. (**A**) fibers of the cellulose/polyester fabric coated with ECM. (**B**) 10,000x magnification of ECM on fabric, showing copper agglomerates in the 3-7 micrometer range.

**Fig 3:**
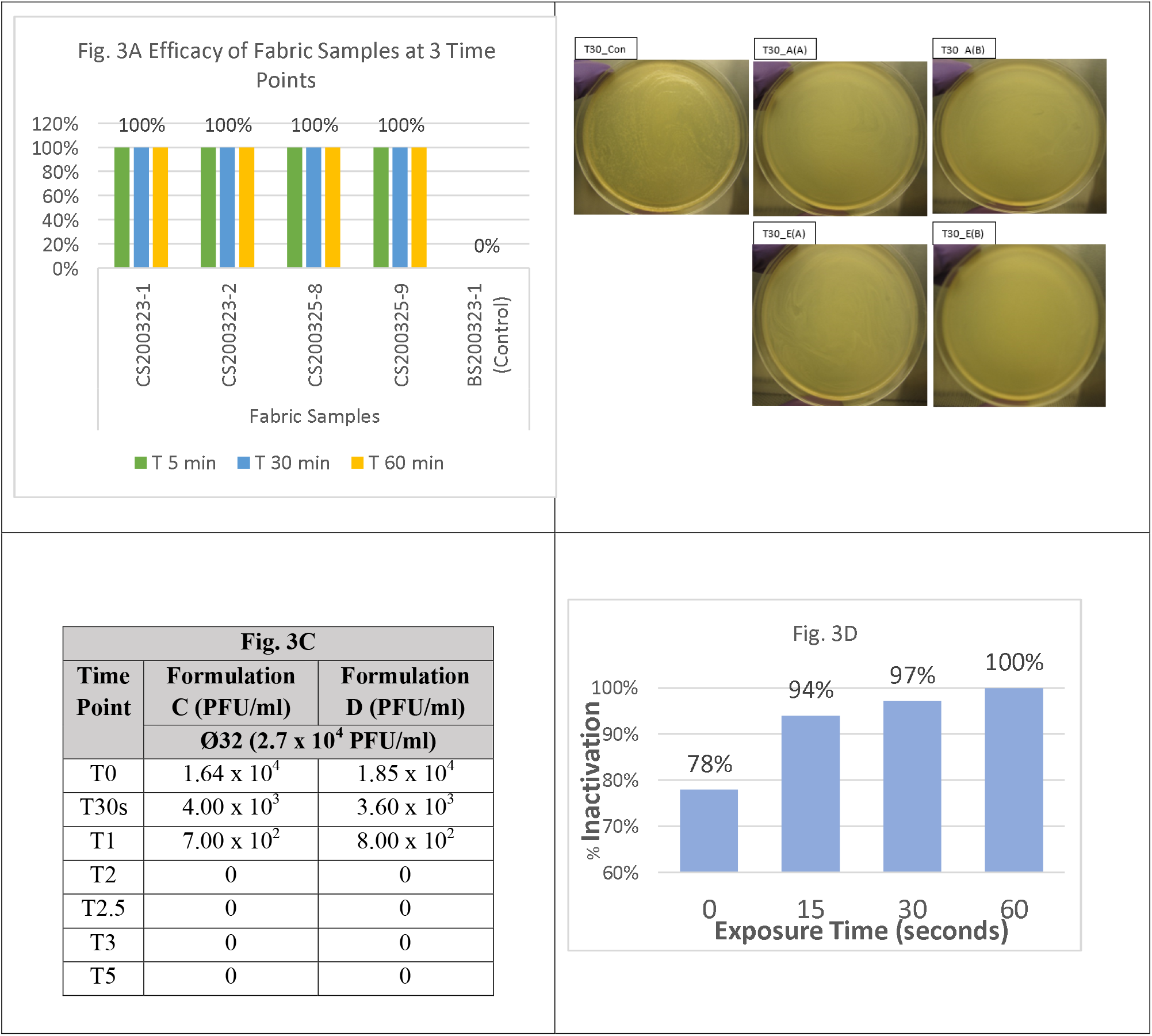
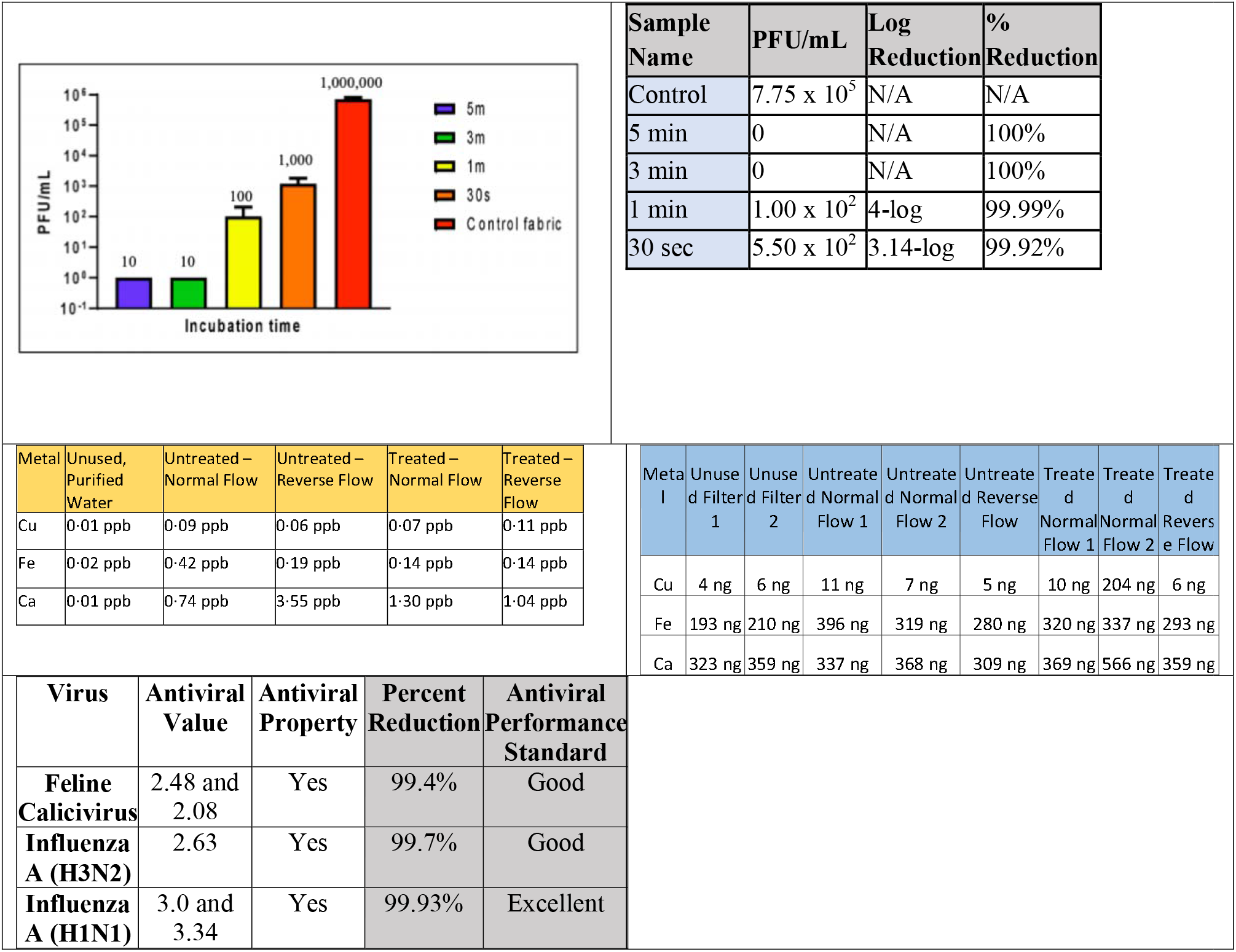
Results from antimicrobial tests of aCu. From left to right, **A** viral kill rate after 5, 30, and 60 minutes, compared to inactive control (4 active samples and 1 control per duration). **B** 100% kill rate of bacteriophage after 30 min on aCu-treat rayon/polyester fabric. **C** deactivation of 2·7*10^4^ (a) and 2·7*10^5^ (b) PFU/ml viral titer by 2 formulations (C&D) of aCu. **D** Deactivation of bacteriophage on 2-year-old aCu-coated aluminum substrate over short duration. **E (a, b)** Deactivation of SARS-CoV-2 on aCu-coated fabric (graph and data chart). **F** and **G** Filter metal testing from collected water (**F**) and ICP-MS (**G**) after shedding/particulate testing. Fe indicates traces of stainless steel – stainless steel sample holders were used. Ca indicates environmental contamination/sample hygiene. Spike of copper at treated normal flow 2 is commensurate with other increases, indicating environmental sample contamination. **H** Results of antiviral testing against feline calicivirus and influenza A (H3N2 & H1N1).

**Fig 4:**
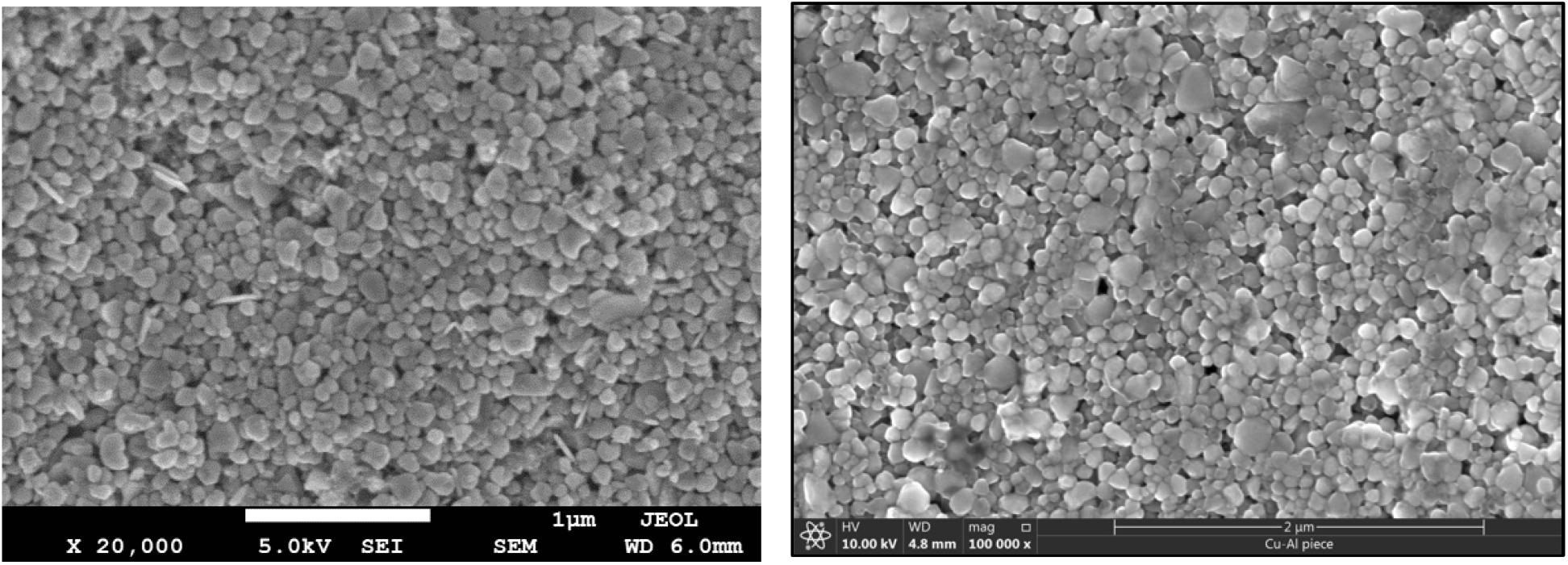
SEM image of two-year-old copper coating layer on Al substrate. **A** after initial coating in 2018; **B** 2 years later. Both show very similar agglomerated particle morphology, shape and size (images have been sized to reflect identical scale).

**Fig 5.**
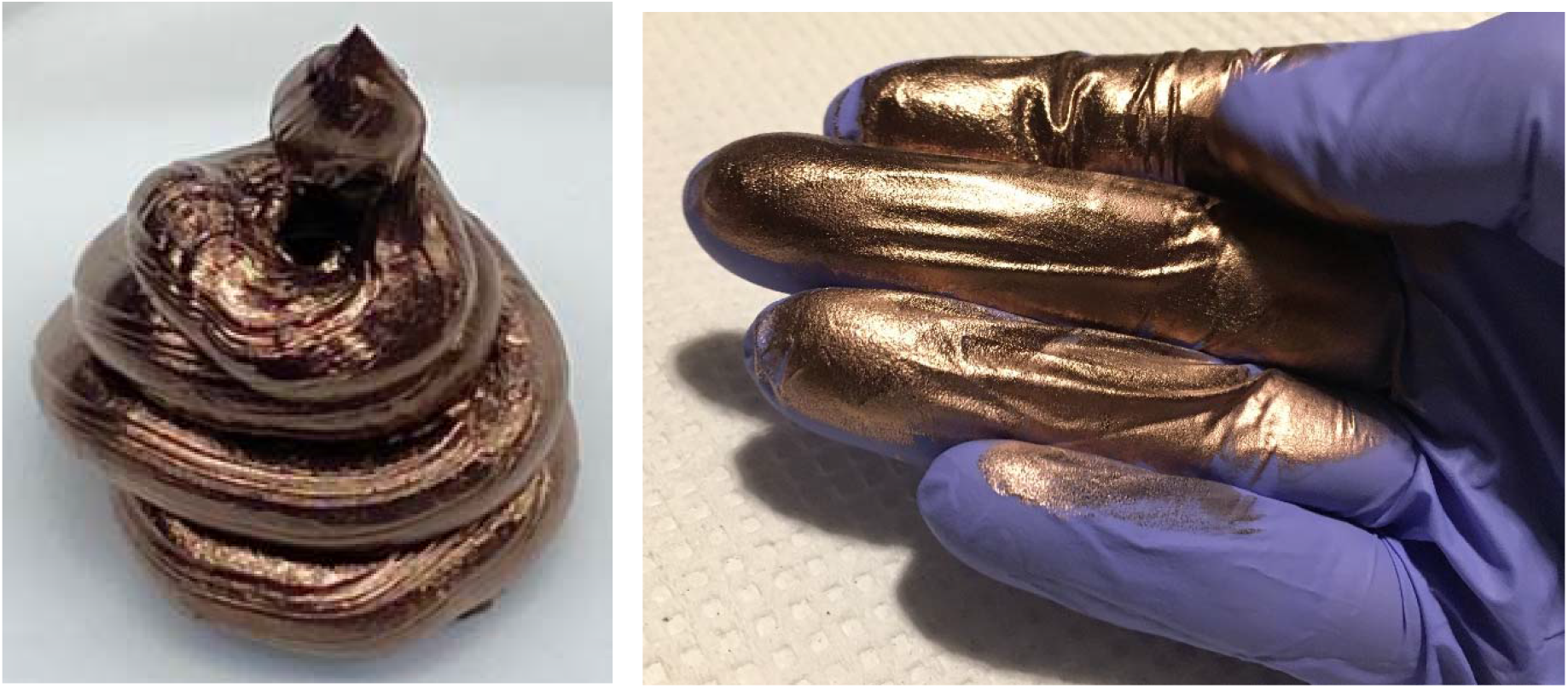
Images of ECM after being dispensed from a tube. The dense thixotropic copper gel is paste-like and stable. It behaves like a typical non-Newtonian “liquid”. A layer of amine surfactants exerts strong cohesion via hydrophobic interactions and organic end-chain entanglement. It can be spread very thin with a lotion-like texture.

We set out to quantify ECM’s activity against a host of viruses and bacteria with the eventual goal of testing against the SARS-CoV-2 virus. We formulated water-based paints to coat fabrics and porous filter materials that could be used in face masks and air purification units such as HVAC systems in airplanes, hospitals or public transportation. For clarity, we will refer to the antimicrobial applications of ECM as ActiveCopper (aCu). The Environmental Protection Agency (EPA) outlines a set of tests to measure the efficacy, longevity, and durability of copper alloys as antimicrobials. In each instance our antimicrobial tests followed the EPA protocols. We made the decision to lengthen and shorten test windows to increase the demand placed on our material. In practice, this meant shortening the pathogen-aCu contact time to truly determine how quickly the antimicrobial action took place and lengthening the duration of multi-day, multi-inoculation tests to gauge how long aCu could be expected to remain active.

## Materials & Methods

### Synthesis of ECM

A 3% solution of CuCl_2_ (3 L of 0.5M) in glyme was heated to 40 °C under a nitrogen atmosphere and 1.4 L of a 3 M NaBH_4_ solution (1.4 M equivalent) in glyme added as the reducing agent. The reaction mixture was continuously stirred for another 20 min until a dark red-brown slurry has formed and then cooled to room temperature. The solids were isolated by centrifugation and then washed with 7 L of water to remove (dissolve) the NaCl that formed during the reduction step and centrifuged a second time to isolate the ECM. The reaction yields about 100 g of product.

### Antimicrobial Testing

#### Modified International Standard Organization ISO Test 18184:2019 against Salmonella enterica-specific bacteriophage Ø32 4/1/2020

The following five tests were conducted by Dr. Mina Izadjoo and Dr. Hosan Kim at Integrated Pharma Services in Frederick, MD. Four aCu-coated samples (CS200323-1, CS200323-2, CS200325-8, CS200325-9) and one control (BS200323-1) (A(A), A(B), E(A), E(B), Cont.) were used in this initial screen of activity against *Salmonella enterica*-specific bacteriophage Ø32. The samples were plated and administered a 5×10^4^ PFU challenge of bacteriophage Ø32. After the predetermined incubation period of 5, 30, or 60 min, 0.2 ml Dey-Engley neutralizing broth (Becton, Dickinson and Company, Franklin Lakes, NJ) was applied to wash the bacteriophages. The samples were removed as quickly as possible to avoid additional contact time. The viral suspension was serially diluted with phosphate buffered saline (PBS, Sigma Aldrich, St. Louis, MO). 0.2 ml of the resultant phage solution was administered to individual tryptic soy agar (TSA, MP Biomedicals, LLC, Solon, OH) plates along with 0.2 ml of the overnight NR-170 culture. 5 ml of 0.7% top agar was added and swirled to adequately mix. TSA plates were incubated at 37 □ for 18 hours before being examined for plaques. 100% virus killing would be indicated by no plaques.

This same procedure was repeated with aCu on rayon/polyester (70/30) fabric samples (BS200401-1 Control, CS200401-1, −2 Active).

#### Modified (Lengthened) EPA Long-Term Test

Twenty cellulose/polyester (45/55) fabric samples cut from batch CS200416-3 (active) and BS200416-1 (control) were used in this trial. Ten were untreated and used as controls, ten were coated with 3 front and 2 back applications of aCu. Samples were challenged with a 2.7 × 10^7^ PFU/ml titer of *Salmonella enterica*-specific bacteriophage Ø32. The samples were tested sequentially; all samples received inoculation on T1, 4/21/20, followed by a 24-hour incubation period. Sample 1 received a single inoculation (4/21), sample 2 received two inoculations (4/21, 4/22), sample 3 received three inoculations (4/21, 4/22, 4/23) and so on up to sample 10, which received ten inoculations (4/21-5/4) over the 14-day period. Inoculations were not administered on Sat/Sun (4/25, 4/26 & 5/2, 5/3). Once the incubation period had elapsed (either 24 hours on weekdays or 72 hours on weekends) the virus was recovered with 1 ml of Dey-Engley neutralizing broth followed by vortexing and centrifugation. The resultant solution was then mixed with *Salmonella enterica* NR-170 culture and 0.7% top agar poured onto a TSA plate. After an 18-hour incubation period, plates were examined for presence of viral plaques. The complete absence of plaques indicated that aCu had achieved a 100% kill rate. The control fabrics were administered the same protocol and displayed prolific plaques at each time point, indicating otherwise hospitable conditions for viral growth.

#### Modified (Shortened) Ultra-Rapid EPA Efficacy Test

Cellulose/polyester (45/55) fabric samples from batches BS200416-2 and CS200416-3 coated with aCu were used in this short-term exposure test. Samples from BS200416-1 were used as a blank control. aCu coated fabric samples were placed on empty petri dishes and spotted with 0.2 ml of the *Salmonella enterica*-specific bacteriophage Ø32 titer in the center of the fabric. This application delivered 2.7 × 10^7^ PFU/ml of bacteriophage to the sample. After inoculation the samples were incubated at room temperature for T0, T30s, T1, T2, T2.5, T3, and T5 minutes. After the designated time had elapsed, 2 ml of Dey-Engley neutralizing solution was added to wash the bacteriophages. The samples were removed as quickly as possible to avoid additional contact time. The viral suspension was diluted with phosphate buffered saline (PBS). 0.2 ml of the resultant phage solution was administered to respective tryptic soy agar (TSA) plates along with 0.2 ml of the overnight NR-170 culture. 5 ml of 0.7% top agar was added and swirled to adequately mix. TSA plates were incubated at 37 L for 18 hours before being examined for plaques. 100% virus killing would be indicated by no plaques, however further growth of host bacteria would continue.

#### Modified (Lengthened) EPA Longevity Test

A 2×2 cm, 3 mm thick piece of aluminum was placed on an empty plate. 100 µl of *Salmonella enterica* specific bacteriophage Ø32 phage titer (∼5 × 10^4^ PFU/ml) was spotted on the center of the aluminum sample. The sample was incubated at room temperature (37 L) for the designated time periods before the addition of 1 ml of Dey-Engley neutralizing broth. The aluminum was promptly washed by wiping with 70% isopropyl alcohol (IPA, Decon Labs, Inc, King of Prussia, PA), followed by light wiping with fabric swaths (CS200416-3 or CS200416-4 fabric samples). The viral suspension was diluted with PBS. 0.1 ml of the resultant solution was spotted on a TSA plate with the addition of 0.1 ml of *Salmonella enterica* NR-170 overnight culture. 5 ml of top agar (0.7%) was added and swirled to mix adequately. The plates were incubated at 37 L overnight before undergoing examination for viral plaques. Absence of plaques indicates 100% kill of bacteriophage

#### Determination of Antiviral Activity of Textile Products based on International Standard Organization (ISO) Test Method 18184:2019

All test specimens were prepared in sterilized, capped vials. 2×2 cm pieces of cellulose/polyester (45/55) fabric were used. Untreated control specimens came from batch BS200427-1, aCu treated active specimens came from batch CS200427-1. S a m p l e s were sterilized via ultraviolet radiation overnight and stored in sterilized vials prior to testing. 0.2 ml of the respective virus suspensions were deposited onto each specimen and caps were tightly closed. The test samples were incubated for 2 hours at room temperature. The blank control samples were immediately treated (T0) with 20 ml of wash solution, SCDLP (Casein Peptone Lecithin Polysorbate Broth with Tween 80) medium. The aCu-coated specimens were treated with 20 ml SCDLP medium after a 2-hour contact time. The vials were closed and agitated by vortexing for 5 seconds repeated 5 times to wash out the virus. Results from antiviral tests of treated textile were compared to results of control (untreated textile).

The wash out suspension from both the control and treated samples and the virus control suspension was serially diluted in Essential Minimum Essential Medium with 1% Penicillin/Streptomycin antibiotics. The Human Influenza A H1N1, Human Influenza A H3N2 and Calicivirus VR-782 viral suspensions were diluted with PBS to concentrations of 10^7^ pfu/ml from the original titer.

For antiviral analysis, host cells to be used for the plaque titer determination assay were prepared in tissue culture 6-well plates and incubated at 37 °C with 5% CO_2_ for up to 2 days. Host cells for the TCID_50_ cytopathic effect titer determination assay prepared in tissue culture 96-well plates and incubated at 37 °C with 5% CO_2_ for 7 days. Each virus has a specific host cell: MDCK Cell ATCC CL-34 is used for H1N1 and H3N2 and CRFK Cell ATCC CL-94 is used for Calicivirus.

1. For TCID_50_ Determination – Behrens and Karber Method was used
  1. Y=X ×10^a^
  2. a = ∑p – 0.5

#### Antiviral Efficacy Test Against SARS-CoV-2

SARS-CoV-2 testing was conducted at the Biomedical Research Laboratory at George Mason University in Manassas, Virginia by Dr. Kylene Kehn-Hall and associates in a BSL-3 laboratory. The test protocol was developed by Dr. Kehn-Hall. aCu-coated and blank control fabric samples from batches CS200429-1 and BS200429-1 respectively were cut into 22 mm diameter disks and added to individual wells of a 12-well plate. 1 ml of Dulbecco’s Modified Eagle Medium (DMEM) (VWR, Radnor, PA) supplemented with 10% FBS (VWR Cat # 97068-085), 1% L-glutamine (VWR Cat # 45000-676), and 1% penicillin/streptomycin (VWR Cat # 45000-652) was placed in each well and samples were incubated for 0, 0.5, 1, 3, or 5 minutes. Following incubation, the cell culture media and CellTiter Glo (Promega, Madison, WI) was removed with a pipette. To obtain as much liquid as possible, the fabric samples were placed in a spin-basket within a microcentrifuge tube. The tubes were centrifuged at 10,000 g for 5 min (Microfuge 16, Beckman Coulter, Indianapolis, IN) and the collected media was combined with the previously pipetted media.

The combined sample was transferred evenly into 8 wells within a 96-well plate. Each of the 8 wells contained Vero cells (ATCC Cat CCL-81). The 96-well plate was incubated in a CO_2_ incubator (5%) at 37 °C overnight; luminescence was measured at 18 hours post-treatment with untreated samples included as a control (GloMax Discovery, Promega, Madison, WI). Data was normalized to the untreated control and presented as the average (N=3) with standard deviations.

Geometric Mean of Test Carrier:

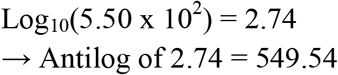

Average of Control Carrier:

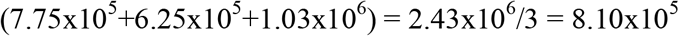

Geometric Mean of Control Carrier:

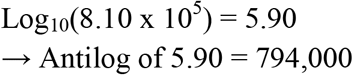

% reduction:

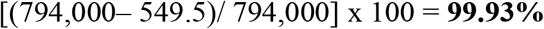

Recovery Log_10_ Difference:

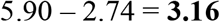

#### Copper Shedding/Breathing Simulation Test

Samples treated with various coats of aCu (BS200408-1 control, CS200401-2, −3, −4, −5, −6 active, Table S1) were individually loaded into a 25 mm stainless steel filter holder (Millipore Sigma, Burlington, MA). Purified dry N_2_ was blown through the filter holder for 15 min at a rate of 4.1 slpm based on estimates of the peak air flow rate experienced by a typical respirator mask during normal breathing. Nitrogen was administered in both the “forward” and “reverse” directions to simulate inhalation/exhalation, and collect shedding activity data from both sides. Optical particle counting using bin sizes of 0.2, 0.3, 0.5, and 1.0 µm was conducted using an Apex R02 Particle Counter (Lighthouse, Fremont, CA) on the N_2_ after passing through the aCu-treated fabric. Condensation particle counting was conducted between an inclusive range of 8 nm-3 µm. A water bubbler trap was used to collect any shed materials from the exhausted N_2_. Inductively coupled plasma mass spectrometry (ICP-MS) was conducted to analyze the water trapping solutions from the flow-through. Filtered N_2_ was captured by a 47 mm filter trap (Millipore Sigma, Burlington, MA) and 0.1 µm Track-Etch filters. These filters were used to look for the evidence of shed metal using SEM-EDS. Samples were submitted to Millipore Imaging Lab in Bedford, MA for imaging.

## Results

As an initial rapid efficacy pre-screen, we tested aCu against *Salmonella enterica*-specific bacteriophage Ø32 with exposure times of 5, 30, and 60 minutes. All active samples showed 100% viral kill rates after just 5 minutes; no plaques remained to be counted (Fig. 3A). To ensure that these dramatic results were accurate, this test was duplicated on a different rayon/polyester (70/30% w/w) fabric blend where it yielded the same result (Fig. 3B).

To determine how quickly aCu acts against microbes, we opted to further decrease the testing time increments to as short as 30 sec and test the kill rate upon initial contact. Cellulose/polyester (45/55) samples were sprayed with two variations of aCu paint for the “ultra-rapid” efficacy test: C4 contained paint alone, D4 included CuCl_2_ as an activator. Viral challenges of *Salmonella* bacteriophage were administered to each of the samples. The remaining phages were measured at seven intervals (T0, T30s, T1, T2, T2.5, T3, T5 min). At T0, reductions of 15,000 PFU/ml (C4) and 62,000 PFU/ml (D4), 6% and 23% respectively, had already taken place (Fig. 3C). By T2, the D4 sample had achieved a 100% kill rate of both phage concentrations (Fig. 3C). Sample C4 achieved 100% kill rates of the 2·7×10^4^ PFU/ml concentration by T2 and the 2·7×10^5^ PFU/ml concentration by T3 (Fig. 3C). This compares very favorably to 70% ethanol, the disinfectant regarded as the standard, which achieved a 3 log_10_ reduction of SARS-CoV-2 after 30 seconds,^7^ and a 2 log_10_ reduction of feline calicivirus after one minute.^8^

Our subsequent test measured performance over an extended period of use to address the durability requirements established by the Environmental Protection Agency (EPA) for the registration of copper alloys as antimicrobial agents. Regulations stipulate that compounds must be able to continuously reduce a recurrent bacterial load,^9^ which would be expected in a hospital setting or on surfaces in public transportation.

Over 14 days, aCu’s sustained antimicrobial efficacy was tested according to the EPA protocol, which calls for repeated inoculations over the course of 21 hours.^9^ We lengthened the timeline to better gauge the longevity we could expect from copper-coated fabrics (fig. S2) and applied a very large viral load of 2·7×10^7^ PFU/ml to simulate high touch-traffic scenarios. The first extended exposure test was planned for 10 days anticipating some performance degradation. aCu-coated cellulose/polyester (45/55) fabric was treated with a daily titer of *Salmonella* bacteriophage Ø32 (2·7×10^7^ PFU/ml). After 10 days, antimicrobial capacity showed no degradation whatsoever. We opted to extend the test to 14 days. After 14 days the aCu-coated sample had maintained its antiviral capacity (fig. S2). The material was as active against phages on day 14 as on day 1. The results of this test demonstrate the high efficacy of aCu even when subjected to the frequent and intense pathogen exposure that is a fixture of health care settings.

We wanted to understand what kind of durability and longevity we could expect from solid surfaces coated with a layer of aCu paint. A 2×2 cm, 3 mm thick piece of aluminum that had been fully coated with aCu paint in 2018 stored indoors in ambient conditions with full exposure to seasonal temperature swings was used for the viral challenge test. By T15 s, 94% of the phages had been killed, by T30 s 97%, and by T60 s fully 100% of the phages had been killed, leaving no viral plaques to count (Fig. 3D). SEM analysis revealed that the copper material had not changed its morphology over the two years, indicating excellent shelf-stability (fig. S4a). From these data we learned that aCu maintains vigorous antimicrobial activity independent of its substrate; textiles and hard surfaces are both viable mediums.

Encouraged by aCu’s durability and fast action, we conducted efficacy testing based on ISO test #18184:2019 – Determination of Antiviral Activity of Textile Products. aCu was applied to cellulose/polyester fabric with additional CuCl_2_ as an activator. Samples were challenged with human influenza A (H1N1 & H3N2) and feline calicivirus VR-782. Viral suspensions were deposited onto the fabric samples and after two hours (shortest allowable with ISO) of incubation the aCu textile had reduced the concentrations of all three viruses by more than 99% (Fig. 3H, Table S6b). H1N1 was reduced by 99·93%, H3N2 by 99·77%, and feline calicivirus by 99·42%. It is encouraging to yield favorable results against both enveloped (influenza) and non-enveloped (calicivirus) viruses.

These preliminary investigations culminated in a test against the SARS-CoV-2 virus to determine the suitability of aCu-treated face masks to prevent virus transmission. The cellulose/polyester (45/55) fabric was coated with aCu and adhesive. A viral load of approximately 10^6^ PFU/ml was applied to the fabric; after 30 seconds up to 99·9% of the virus had been killed (Fig. 3Ea, b, Table S5). The results are represented in the equation below.

Geometric Mean of Test Carrier:

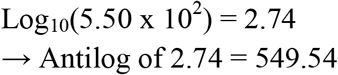

Average of Control Carrier:

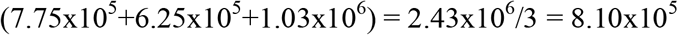

Geometric Mean of Control Carrier:

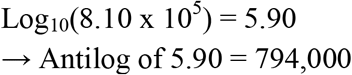

% reduction:

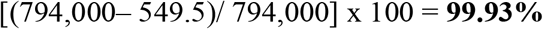

Recovery Log_10_ Difference:

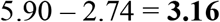

This dramatic efficacy makes aCu a very promising candidate for the development of self-cleaning PPE.

### Copper in medicine and mammalian compatibility

To ensure that no harmful respiratory exposure occurs from using an aCu-coated insert in a face mask, shedding tests were conducted under simulated breathing conditions. aCu-coated fabric and blank controls were individually loaded into a sample holder and a flow of nitrogen was administered at rates commensurate with breathing. Two different flow rates were tested, 4·1 slpm and 20 slpm, to simulate normal breathing and more intense breathing during physical activity. A full battery of tests, including optical particle monitoring, condensation particle counting, filter trapping, and water trapping were employed to assess the exhausted N_2_ for shed particles (Fig. 3F, G). It should be noted that these methods detect all potentially shed particles, including copper, fabric fibers, dust, or others. “Total particles” refers to all particles that lifted off the fabric under N_2_ flow. SEM/EDS analysis of the filter traps did not detect any copper (fig. S4). Metal levels in the water trapped from both treated and blank control materials were equivalent; no Cu was detected (Fig. 3F). Optical particle counter tests found total particle counts to be <0·1 ng in each 15-minute test window (fig. S3). These particles were organic in nature and looked like loose fabric material. Hypothetically, if all of the identified particles were copper, it would equate to a maximum shedding rate of 0·1 ng Cu per 15-minute period. This would translate to a maximal copper “dose” of approximately 2 ng/hour for a wearer. This would be significantly below OSHA’s maximum Permissible Exposure Limit (PEL) of 1mg/m^3^ for copper dust in the workplace.^10^ The simulated breathing data offers reassurance that aCu-coated fabrics are safe for frequent human use and proximity to the respiratory system.

Our results far exceed the benchmarks established by the EPA for speed of action as well as material longevity and durability. The favorable results against multiple virus configurations (enveloped and non-enveloped) suggest that aCu is broadly lethal against pathogens and is therefore unaffected by the possibility of future mutations. Enveloped viruses, like coronaviruses and influenzas, are generally regarded as less hardy than non-enveloped viruses.^11^ aCu’s non-selectivity coupled with its rapid deactivation window suggests viability as an additive to healthcare tools like privacy curtains, door handles and guard rails.

As with any material that contacts skin or the respiratory system, safety is of utmost concern. A protective item must not introduce new health threats while protecting the user. To this end, it’s pertinent to highlight copper’s long and safe history in medical and industrial materials.^12, 13^ Ancient civilizations recorded medical copper use as early as 2200 B.C., and a more scientific understanding was established when a doctor realized that copper miners and smelters enjoyed distinct immunity from Paris’s 1832 cholera outbreak.^*4*^ Reliance on copper continued to grow until it was displaced by penicillin in 1932. More recently, copper has been incorporated into medical devices,^14^ dental cement,^15, 16^ and dilute CuCl_2_ IV solutions to treat copper deficiencies in patients.^17^ Copper’s bioactivity is relevant in both the synthesis and maintenance of skin proteins; it is used in treating skin conditions.^18^ Extended dermal exposure to copper has been shown to be well-tolerated.^19^

Copper plays an essential role in human health. It is involved in angiogenesis,^20^ transcription factor regulation, bone development, and various other physiological processes.^21^ Deficiencies in bodily copper can cause anemia, neutropenia, and neurological disorders.^21, 22^ Because it is so well integrated into human biology, robust mechanisms exist to regulate copper levels in the body.^23^ Homeostatic maintenance of copper occurs via intestinal absorption and release is mediated by the liver into bile.^24^ Small stores of copper are kept in the body, approximately 0·5 mg per pound of body weight in the average adult.^25^

## Discussion

When the EPA recognized the antimicrobial efficacy of copper alloys in February, 2008, copper’s biocompatibility contributed to their determination that “these products pose no risks to public health; copper products have been in use for centuries, and we know of no harm from such use”.^26^ EPA recognition allows copper manufacturers to claim that approved copper products “kill 99·9% of bacteria within two hours”.^26^ The two-hour guarantee of bacterial deactivation is accompanied by an EPA mandate to maintain rigorous disinfecting regiments because copper that works on this timescale could still transmit pathogens. Over 500 copper alloys are registered with the EPA and deemed to have no “unreasonable adverse effects” under the U.S. Federal Insecticide, Fungicide, and Rodenticide Act.^26^

Because our testing indicated that aCu has the potential to function as personal protective equipment in a medical setting, it is necessary to consider the precedent set by other medical copper products prior to the development of aCu for safety and indicators of need within the field. We considered the guidance from the Food and Drug Administration (FDA). The FDA has listed fine copper powder (95% w/w Cu) as exempt from certification, deeming it safe for use as a color additive in cosmetics. This qualification was extended to products intended for use specifically in the eye area.^27^ The FDA’s relaxed position on copper provided further reassurance that our material would create no health issues for the people using it. FDA approval has been sought and granted for other medical copper products, establishing a precedent for copper’s enduring modern utility. FDA approval was granted for a breathable face mask in 2016^*28*^ that protects against bacteria and influenza viruses. The mask material was proven to kill 99% of tested bacteria after one hour of contact and inactivate 99·9% of test influenza viruses after five minutes of contact.^28^

The antimicrobial technology at play applies fine zeolite infused with Ag^+^ and Cu^+^ ions to the non-woven polyester fibers of a mask. Their application method eliminated off-gassing or leaching of the antimicrobial agent, which we have accomplished with aCu (fig. S3, S4). To securely fix aCu paint to the intended material, we have adopted the practice of gluing (bonding) aCu to the core of the fabrics. aCu’s protective lipid-type layer both protects and stabilizes the high surface area of aCu, further enabling strong cohesion of the micron-size agglomerates (Fig. 1). Our highly active copper material is differentiated from related precursors by its ability to kill viruses and bacteria in one minute or less.

We have speculated on the mechanisms responsible for aCu’s pathogen-agnostic eradication of microbes and have been informed extensively by the work of other researchers. Research on inorganic aerogels, xerogels and the like has shown that they exhibit high surface areas due to their fine pore structure; they are strongly attracted to one another and other surfaces via hydrophobic interactions or hydrophilic hydrogen bonding and chain entanglement.^29^ This can be enhanced by placing long-chain organic surfactants at their surfaces, forming a reverse micelle-type structure. We did exactly that with our aCu and designed a surfactant monolayer that maximizes attraction and mimics a lipid cell membrane. A virus’s small size (20-300 nm) combined with its commensurate surface energy to aCu results in mutual attraction. When they stick together, the protective lipid envelope and the protein capsid of the virus is readily attacked by aCu. The copper oxidizes first to Cu^+1^ and then more rapidly to Cu^+2^ with the formation of very aggressive OH* radicals in the presence of moisture and air as well as fatty acid radicals from the oxidation of the protective lipid layer:

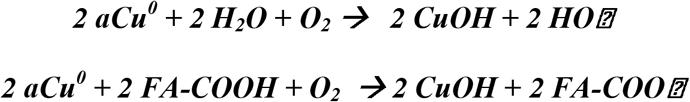

This process causes the lipid layer to disintegrate and the protein capsid to break open .^30, 31^ The destruction of the capsid was identified by Dr. Sarah Warnes, *et al*. in 2015 using transmission electron microscopy (TEM).^32^ Viral replication is halted as Cu^2+^ ions, stabilized by their amine surfactants, cleave and cross-link RNA/DNA through their affinity for nucleotides.^33^ This mechanism is analogous to the DNA binding exhibited by anti-tumor *cis*-platinum compounds.^34^ Both reactive oxygen species and ionic interactions have been shown to facilitate viral deactivation. Due to the unprecedented speed and activity of aCu, more testing is necessary to elucidate the exact order of operations.

Action against bacteria begins as their unique cell wall degrades.^35^ Copper ions inhibit the transpeptidases responsible for cross-linking the peptidoglycan wall, compromising its structural integrity. Bacteriolysis is furthered by copper’s interruption of the necessary balance between peptidoglycan-synthesizing enzymes and autolysin.^35^ The destruction of such a fundamental piece of the bacterial framework in both Gram negative and positive strains indicates that aCu’s efficacy would not be hindered by bacterial mutation. Additionally, antimicrobial copper experiments have been successfully conducted against a number of antibiotic-resistant strains. Reactive oxygen species generated by Cu^2+^’s interactions with the cell wall and subsequent permeation into the membrane may harm bacteria via cytoplasmic toxicity.^36^ aCu does not act as a mutagen, as antibiotics can,^37^ indicating that it is not liable to produce mutant bacterial strains and contribute to antibiotic resistance.^38^

Around the world, people need to return to life, work, and family while protecting their health. A growing number of people who experienced flu-like Covid-19 and were believed to have recovered are reporting persistent fatigue, gastrointestinal, and cognitive issues that outlast their original symptoms.^39^ In some cases, these long-term impacts manifest as cardiac dysfunction.^40^ Trailing symptoms are not unprecedented with coronaviruses; a study conducted in 2009 on survivors of SARS determined that 40% of respondents struggled with chronic fatigue.^41^ In addition to the detrimental follow-on effects of Covid-19, it is becoming apparent that reinfection may be possible.^42^ The uncertainty that surrounds the long-term consequences of contracting Covid-19 puts additional pressure on people to mitigate their risk, no matter their age.

We have just witnessed the most rapid development of a vaccine in history.^43^ Major logistical hurdles remain before the desired rate of distribution can be reached. At this moment, public distrust in medicine and governing authorities translates to considerable vaccine hesitancy in a multitude of countries.^44–46^ This apprehension will likely hinder the compliance necessary to achieve broad inoculation and subsequent immunity.^47^ The lightning-fast arrival of multiple effective vaccines has been an unequivocal triumph. However, a long-lasting protective mask offers individuals an immediate and reliable defense against COVID-19 while circumventing medical skepticism. It is both reversible and individually administered; a vaccine is not.

Remedies targeted specifically at SARS-CoV-2 are in dire need, but in the long term we need broad, non-selective protection against pathogens. In real time we have watched SARS-CoV-2 jump from humans to large populations of farmed mink in Denmark and now we are seeing a second episode of zoonosis as farm workers become sick with the mink-mutated strain.^48^ Genomic sequencing confirmed that this strain was transmitted from humans to animals and back once more. When given the opportunity to move through a dense concentration of hosts, such as a farm, viruses can mutate into more virulent forms much more quickly than they would in the wild.^49^ A reassortant H1N1 influenza virus with pandemic potential has already been identified in China,^50^ and we can continue to count on the appearance of seasonal illnesses. Due to the lag time in development, a situation is conceivable where the distributed vaccine would miss the currently circulating coronavirus mutant. We posit that masks and other protective pieces that self-sterilize to destroy 100% of pathogens could aid in an expedited return to uninterrupted life.

## Acknowledgments

We have been the very fortunate recipients of strong scientific collaboration focused on putting an end to Covid-19. We would like to thank Dr. Jelena Sepa, Ph.D of EMD Performance Materials Corp. for facilitating our particulate testing conducted by James Nehlsen at Versum/Merck. We would like to thank Gilman Louie of Alsop Louie as well as the Defense Innovation Unit (DIU) of the Department of Defense (DoD) for helping to facilitate the antimicrobial testing of aCu.

## Author contributions

**AAZ**: originally conceived of and developed ActiveCopper, manages all projects related its use. **MI, HK** conducted foundational antimicrobial research **KKH, CW, LP, RF** conducted SARS-CoV-2 research **RLB** wrote the paper and compiled data **RR, AV, KKN, NTN, HTZ** produced and managed testing materials, formulated the copper paste, troubleshot technical materials issues **NA** supervised the project and handled administration and funding acquisition **RMS** handles the commercial manufacturing and scale up of the materials, developed methodologies and supervised the projects.

## Data and materials availability

We are in pre-production phase currently manufacturing 40-50 kg of base ActiveCopper per month; the material is commercially available. Bacteria and viruses used in studies are available commercially from ATCC, ascension numbers are included in Supplementary Materials.

