## Supplementary Materials for "Rapidly self-sterilizing PPE capable of 99.9% SARS-CoV-2 deactivation in 30 seconds"

**This PDF file includes:**

Figs. S1 to S4

Tables S1 to S6

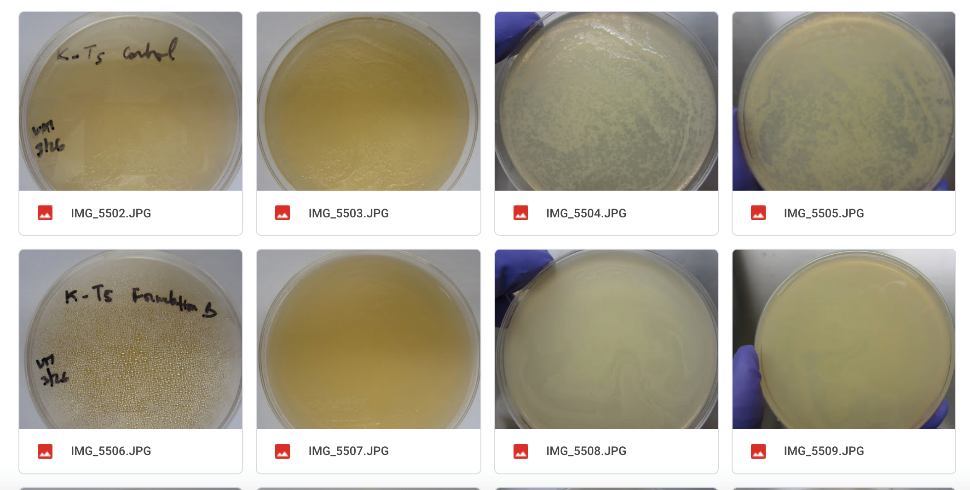

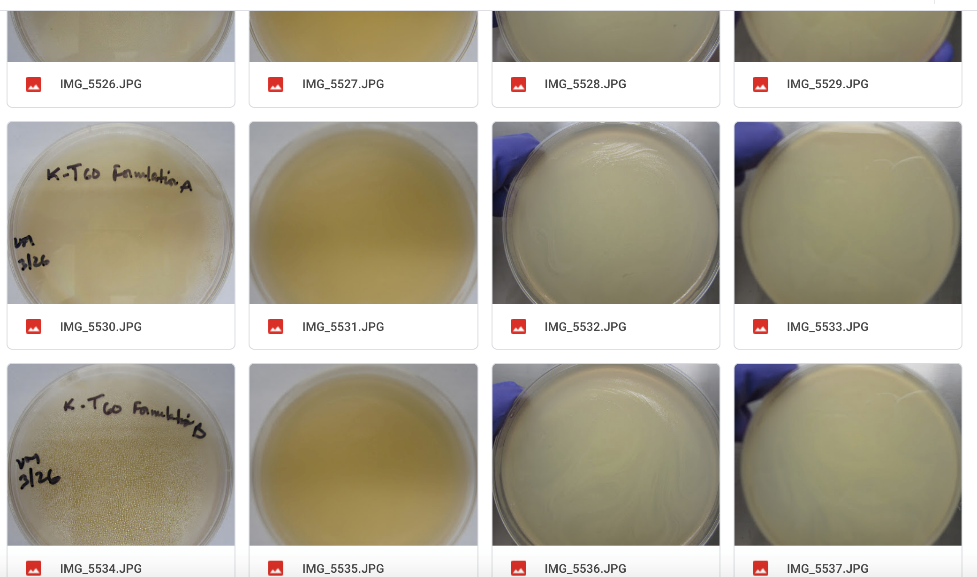

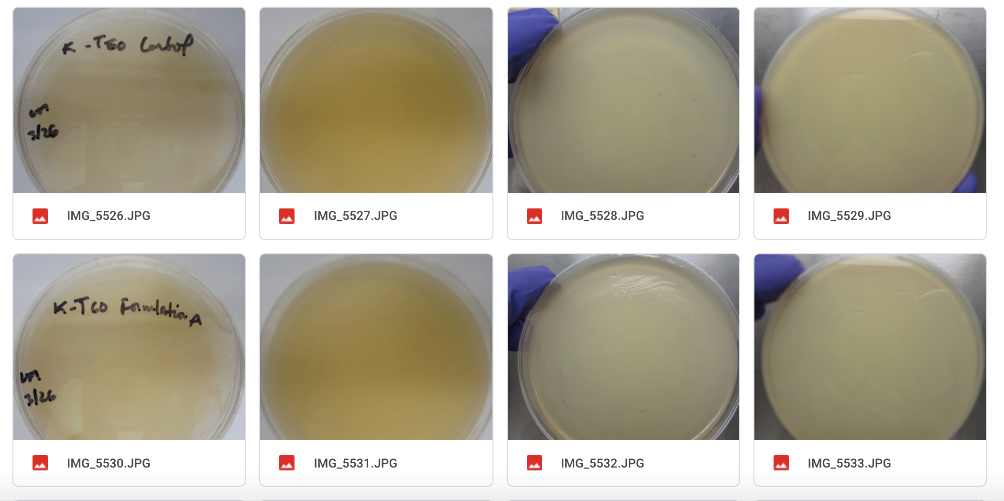

Fig. S1. Images from initial test at T5 & T60 comparing bacteriophage growth between active fabrics and control. A Control & sample B at T5, B Formulation A & B at T60, C Control & Formulation A at T60.

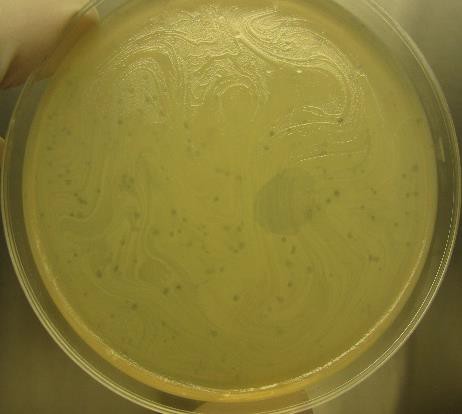

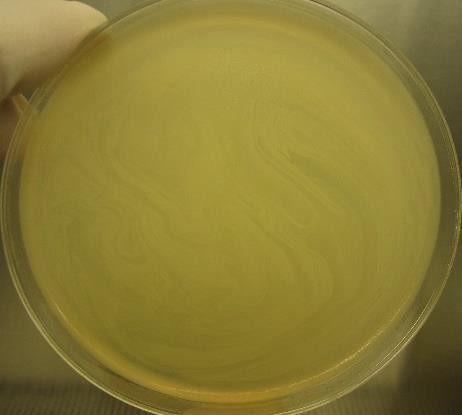

T14-day Antiviral Fabric

T14-day Control Fabric

Fig. S2. Comparison between control and active aCu fabric after 14-day test.

Comparison shows prolific viral plaques on control fabric and no viral plaques on antiviral fabric, indicating 100% kill rate against 107 pfu/ml challenges.

**Comparison of Particles From KuprionTM Samples at Low Flow**

**25mm Disc – 20slpm – Normal flow Direction**

Fig. S3. Comparison of particles from high/low flow rate during breathing simulation test.

CS200408-5 at 15 minutes may be an erroneous read.

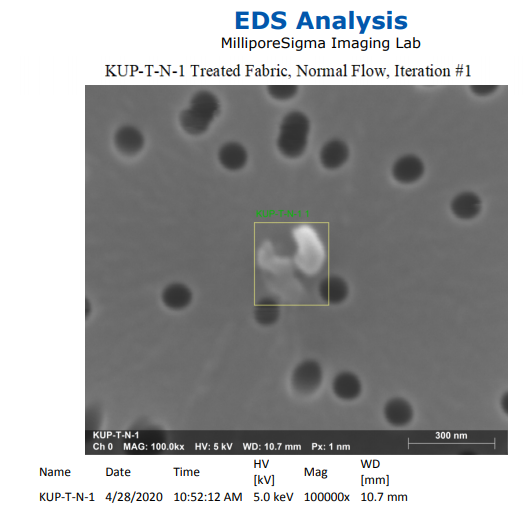

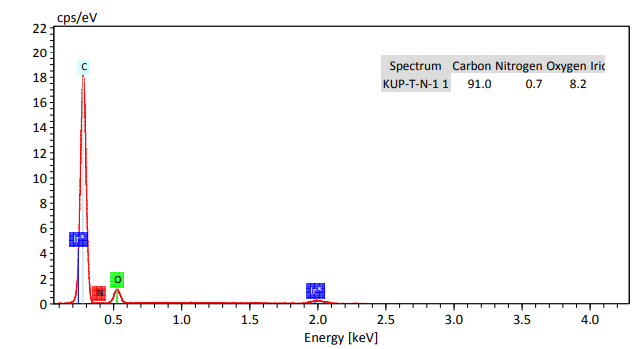

**Fig. S4. SEM-EDS analysis filter trap after flow of nitrogen through treated fabric. Iridium metallization of the samples was used to reduce charging.**

| Sample ID | Description | User | Ship Date | Spray Size | Sample Size | Qty. | Substrate | Spray Dist. | Binder (front) | Cu (front) | Cu (back) | Binder (front) | Binder (back) |
| --- | --- | --- | --- | --- | --- | --- | --- | --- | --- | --- | --- | --- | --- |
| BS200323-1 | Blank Sample | Integrated Pharma Services | 3/23/2020 | 9" x 9" | 2" x 2" | 20 | MicroTek (45/50) | N/A | none | none | none | none | none |
| CS200323-1 | Cu Sample | Integrated Pharma Services | 3/23/2020 | 9" x 9" | 2" x 2" | 20 | MicroTek (45/50) | ~5 | 2x | 3x | 2x | none | none |
| CS200323-2 | Cu Sample | Integrated Pharma Services | 3/23/2020 | 9" x 9" | 2" x 2" | 20 | MicroTek (45/50) | ~5 | 2x | 3x | 2x | none | none |
| CS200325-8 | Cu Sample | Integrated Pharma Services | 3/25/2020 | 9" x 9" | 2" x 2" | 20 | MicroTek (45/50) | ~5 | 2x | **5x** | 2x | none | none |
| CS200325-9 | Cu Sample | Integrated Pharma Services | 3/25/2020 | 9" x 9" | 2" x 2" | 20 | MicroTek (45/50) | ~5 | 2x | 3x | 2x | none | none |
| BS200401-1 | Blank Sample | Integrated Pharma Services | 4/1/2020 | 9" x 9" | 2" x 2" | 20 | U-T Rayon/Poly (70/30)(roll) | N/A | none | none | none | none | none |
| CS200401-1 | Cu Sample | Integrated Pharma Services | 4/1/2020 | 9" x 9" | 2" x 2" | 20 | U-T Rayon/Poly (70/30)(roll) | ~5 | 2x | 3x | none | 1x | none |
| CS200401-2 | Cu Sample | Integrated Pharma Services | 4/1/2020 | 9" x 9" | 2" x 2" | 20 | U-T Rayon/Poly (70/30)(roll) | ~5 | 2x | 1x | none | 1x | none |
| BS200408-1 | Blank Sample | Versum Materials | 4/8/2020 | 12" x 12" | 12" x 12" | 1 | MicroTek (45/50) | N/A | none | none | none | none | none |
| AS200408-2 | Adhesive Sample | Versum Materials | 4/8/2020 | 12" x 12" | 12" x 12" | 1 | MicroTek (45/50) | 3" | 2x | none | none | none | none |
| CS200408-3 | Cu Sample | Versum Materials | 4/8/2020 | 12" x 12" | 12" x 12" | 1 | MicroTek (45/50) | 3" | 2x | 3x | 2x | none | none |
| CS200408-4 | Cu Sample | Versum Materials | 4/8/2020 | 12" x 12" | 12" x 12" | 1 | MicroTek (45/50) | 3" | 2x | 5x | 2x | none | none |
| CS200408-5 | Cu Sample | Versum Materials | 4/8/2020 | 12" x 12" | 12" x 12" | 1 | MicroTek (45/50) | 3" | 2x | 3x | 2x | 1x | none |
| CS200408-6 | Cu Sample | Versum Materials | 4/8/2020 | 12" x 12" | 12" x 12" | 1 | MicroTek (45/50) | N/A | none | 3x | 2x | none | none |
| BS200416-1 | Blank Sample | Integrated Pharma Services | 4/16/2020 | 12" x 12" | 2" x 2" | 25 | MicroTek (45/50) | N/A | none | none | none | none | none |
| BS200416-2 | Cu Sample | Integrated Pharma Services | 4/16/2020 | 12" x 12" | 2" x 2" | 25 | MicroTek (45/50) | 3" | 2x | 3x | 2x | none | none |
| CS200416-3 | Cu Sample | Integrated Pharma Services | 4/16/2020 | 12" x 12" | 2" x 2" | 25 | MicroTek (45/50) | 3" | 2x | 3x | 2x | none | none |
| BS200416-1 | Blank Sample | Integrated Pharma Services | 4/16/2020 | N/A | 2" x 2" | 25 | MicroTek (45/50) | N/A | none | none | none | none | none |
| BS200417-1 | Blank Sample | George Mason University | 4/17/2020 | 12" x 12" | 22mm circle | 10 | MicroTek (45/50) |  | none | none | none | none | none |
| CS200416-2 | Cu Sample | George Mason University | 4/17/2020 | 12" x 12" | 22mm circle | 20 | MicroTek (45/50) | 3" | 2x | 3x | 2x | none | none |
| BS200427-1 | Blank Sample | Integrated Pharma Services | 4/27/2020 | 10" x 11" | 2" x 2" | 25 | U-T Rayon/Poly (70/30)(roll) |  | none | none | none | none | none |
| CS200427-2 | Cu Sample | Integrated Pharma Services | 4/27/2020 | 10" x 11" | 2" x 2" | 20 | U-T Rayon/Poly (70/30)(roll) |  | 2x | 2x | none | 1x | none |
| BS200429-1 | Blank Sample | George Mason University | 4/29/2020 | 11.75" x 11.875" | 22mm circle | 40 | MicroTek (45/50) | N/A | none | none | none | none | None |
| CS200429-1 | Cu Sample | George Mason University | 4/29/2020 | 11.75" x 11.875" | 22mm circle | 90 | MicroTek (45/50) | N/A | 2x | 2x | none | 1x | None |

Table S1. Fabric samples used for testing.

This comprehensive list of samples includes date of manufacture, destination, size, material, coating, and labeling convention.

| **Smpl #** | **Inoculation Date** | | | | | | | | | | **Titer check** | **Incubation time** |
| --- | --- | --- | --- | --- | --- | --- | --- | --- | --- | --- | --- | --- |
|  | **1st** | **2nd** | **3rd** | **4th** | **5th** | **6th** | **7th** | **8th** | **9th** | **10th** |  |  |
| 1 | 4/21(Tue) |  |  |  |  |  |  |  |  |  | 4/23(Thu) | 1-day |
| 2 | 4/21(Tue) | 4/22(Wed) |  |  |  |  |  |  |  |  | 4/24(Fri) | 2-day |
| 3 | 4/21(Tue) | 4/22(Wed) | 4/23(Thu) |  |  |  |  |  |  |  | 4/27(Mon) | 3-day |
| 4 | 4/21(Tue) | 4/22(Wed) | 4/23(Thu) | 4/24 (Fri) |  |  |  |  |  |  | 4/28(Tue) | 6-day |
| 5 | 4/21(Tue) | 4/22(Wed) | 4/23(Thu) | 4/24 (Fri) | 4/27(Mon) |  |  |  |  |  | 4/29(Wed) | 7-day |
| 6 | 4/21(Tue) | 4/22(Wed) | 4/23(Thu) | 4/24 (Fri) | 4/27(Mon) | 4/28(Tue) |  |  |  |  | 4/30(Thu) | 8-day |
| 7 | 4/21(Tue) | 4/22(Wed) | 4/23(Thu) | 4/24 (Fri) | 4/27(Mon) | 4/28(Tue) | 4/29(Wed) |  |  |  | 5/1(Fri) | 9-day |
| 8 | 4/21(Tue) | 4/22(Wed) | 4/23(Thu) | 4/24 (Fri) | 4/27(Mon) | 4/28(Tue) | 4/29(Wed) | 4/30(Thu) |  |  | 5/4(Mon) | 10-day |
| 9 | 4/21(Tue) | 4/22(Wed) | 4/23(Thu) | 4/24 (Fri) | 4/27(Mon) | 4/28(Tue) | 4/29(Wed) | 4/30(Thu) | 5/1(Fri) |  | 5/5(Tue) | 13-day |
| 10 | 4/21(Tue) | 4/22(Wed) | 4/23(Thu) | 4/24 (Fri) | 4/27(Mon) | 4/28(Tue) | 4/29(Wed) | 4/30(Thu) | 5/1(Fri) | 5/4(Mon) | 5/6(Wed) | 14-day |

Table S2. 14-day inoculation schedule for EPA long-term efficacy test.

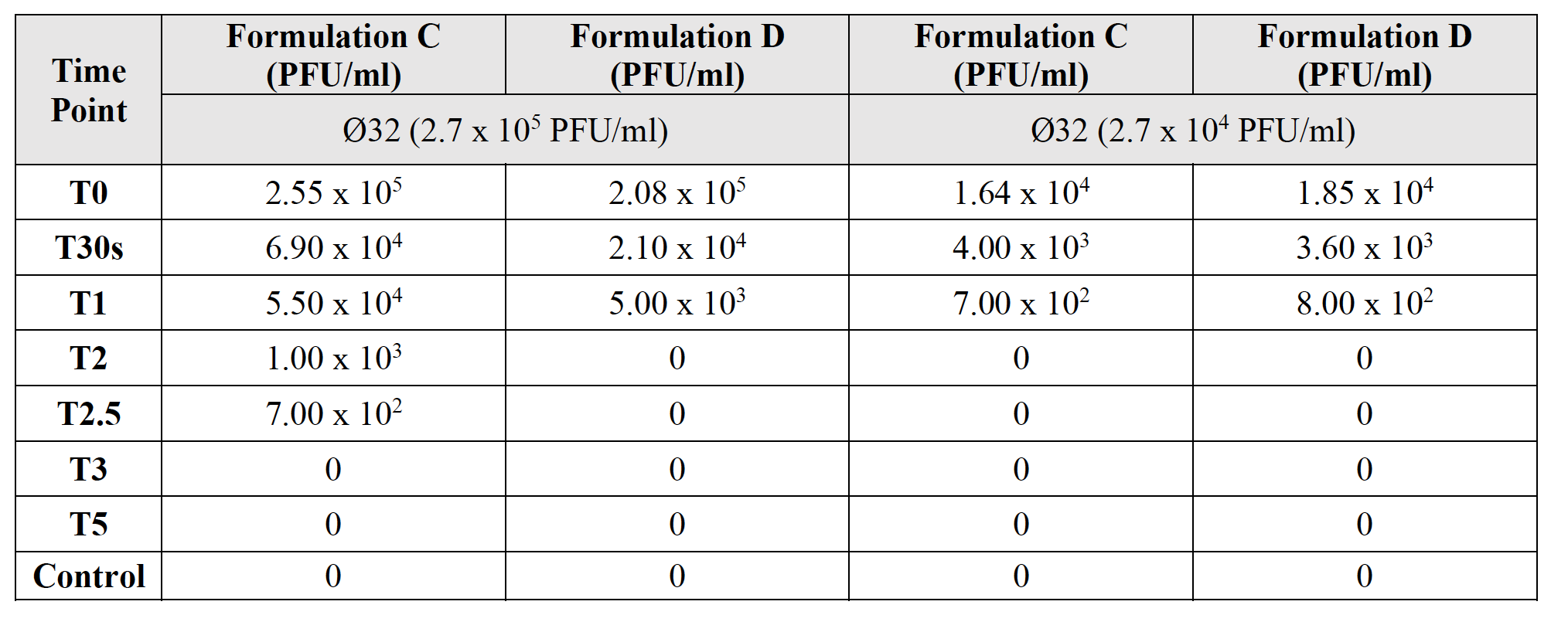

Table S3. Results from ultra-rapid EPA efficacy test.

| **Time Point (sec or**  **min)** | **PFU/ml recovered** |
| --- | --- |
| **T0** | 1.1 x 10^4^ |
| **T15s** | 3.0 x 10^3^ |
| **T30s** | 1.4 x 10^3^ |
| **T1** | 0 |
| **T1.5** | 0 |
| **T2** | 0 |
| **T3** | 0 |
| **Control (PBS)** | 0 |

Table S5. aCu-coated aluminum piece exposure results.

| **Sample Name** | **Replicate** | **PFU/ml** | **Log Reduction** | **% Reduction** |
| --- | --- | --- | --- | --- |
| Control | 1 | 7.75E+05 | 0.12-log | 24% |
|  | 2 | 6.25E+05 | 0.21-log | 38.4% |
|  | 3 | 1.03E+06 | N/A | N/A |
| 5 min | 1 | 0 |  | 100% |
|  | 2 | 0 |  | 100% |
|  | 3 | 0 |  | 100% |
| 3 min | 1 | 0 |  | 100% |
|  | 2 | 0 |  | 100% |
|  | 3 | 0 |  | 100% |
| 1 min | 1 | 1.00E+02 | 3.90-log | 99.9% |
|  | 2 | 0 |  | 100% |
|  | 3 | 2.00E+02 | 3.60-log | 99.9% |
| 30 sec | 1 | 1.30E+03 | 2.79-log | 99.8% |
|  | 2 | 5.50E+02 | 3.16-log | 99.9% |
|  | 3 | 1.68E+03 | 2.68-log | 99.7% |

Table S5. Data from SARS-CoV-2 Deactivation Test. Batch: BS200529-1 & CS200529-3

Formulation ”D”, 0.044g CuCl2, wipe, 3x/2x + 3x 25% adhesive. About 3 mg/in^2^. Average of control PFU = 8.10x10^5^.

| **Virus** | **Antiviral Value** | **Antiviral Property** | **Percent Reduction** |
| --- | --- | --- | --- |
| Feline Calcivirus | 2.48 and 2.08 | Yes | 99.42% |
| Influenza A (H3N2) | 2.63 | Yes | 99.77% |
| Influenza A (H1N1) | 3.0 and 3.34 | Yes | 99.93  % |

| Item | Antiviral Efficacy value (Mv) | Standard |
| --- | --- | --- |
| Tested textile product | 3.0 > Mv > 2.0 | Good Effect |
|  | Mv > 3.0 | Excellent Effect |

Table S6a, b. Antiviral Performance Standard and results of ISO Antiviral Activity Test.
